# Comprehensive Network of Signaling Pathways in Hepatocellular Carcinoma and Network Analysis Reveals PTK2, GSK3β, and β-catenin to be Crucial for Cancer Progression

**DOI:** 10.1101/2025.07.25.666714

**Authors:** Sai Bhavani Gottumukkala, Akilan Malaieaswaran, Anbumathi Palanisamy

## Abstract

Hepatocellular carcinoma (HCC) is a highly aggressive primary liver cancer with significant morbidity and mortality. The development and progression of HCC involves several genetic mutations, alterations in signaling pathways leading to dysregulation of cellular processes. Several investigations have demonstrated the involvement of diverse regulators and their complex signaling cascades in progression of HCC. Understanding these crucial signaling pathways, including receptor tyrosine pathways, PI3K/AKT/mTOR, Wnt/β-catenin, Ras/Raf/MAPK, TGF-β, Hedgehog, JAK/STAT, and Hippo signaling, offers potential avenues for further experimental explorations and system level studies. The present study aims to construct a comprehensive molecular map of the signaling pathways involved in the development and progression of HCC using network editor Cell Designer. The assembled network, consisting of 193 distinct regulators and 362 reactions, provides a holistic representation of HCC associated pathways and adheres to Systems Biology Graphical Notation (SBGN) format. Through subsequent analysis the crucial regulators such as PTK2, GSK3β, β-catenin, and their pivotal roles in driving HCC progression were illuminated. These findings underscore the potential prognostic and clinical significance of regulators influencing the patient outcomes in HCC. Similar workflow can provide further insights and comprehensive understanding of the disease mechanisms.

**Highlights:**

- A detailed molecular interaction map of pathways and regulators involved in HCC was developed.
- To illustrate one of the applications of the map developed, topological analysis using Cytoscape was performed to identify the significant hubs. Subsequently, the prognostic relevance of these identified hubs was evaluated using the transcriptome-based analysis.
- These analysis revealed the crucial roles of regulators like PTK2, GSK3β, and β-catenin offering new prospects for potential prognostic and clinical significance of regulators in influencing HCC.

**Graphical Abstract:** 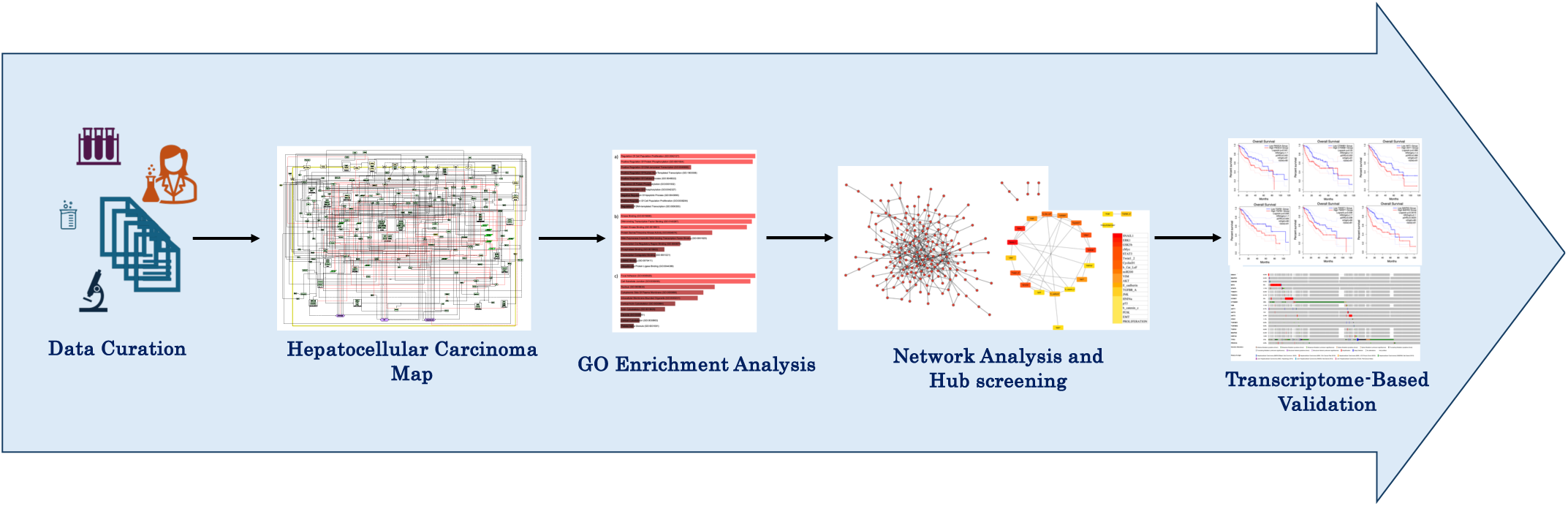

## Introduction

Hepatocellular carcinoma (HCC) or hepatoma is a most common type of primary liver cancer and one of the leading cause of cancer related deaths globally [1, 2]. The most important and independent risk factors include hepatitis B/C virus, persistent alcohol use, autoimmune hepatitis, and various lifestyle choice factors that increase the risk of HCC [3, 4]. Conventional surgical and non-surgical treatment methods were known to be effective. However, these treatments often fall short in effectively curbing the growth, advancement, and metastasis of HCC [5–7]. Recent research underscores the significant interplay between the etiological factors and microenvironment on HCC development and progression [8]. This interplay is crucial in modulating various aspects of hepatocellular carcinoma (HCC), including carcinogenesis, epithelial-mesenchymal transition (EMT), invasion and metastasis [9]. Understanding these influencing factors may yield insights for shaping effective strategies for combinatorial drug targets and treatment strategies.

Molecular pathogenesis of HCC encompasses a complex, multistep process involving several signaling pathways [10, 11]. The activation of specific pathways and the acquisition of a tumor phenotype distinguishes proliferating cells from non-progenitor cells [12]. During the progression of HCC, these pathways undergo dysregulations disrupting the delicate balance between cell proliferation and cell death leading to uncontrolled cell division and metastasis of hepatocytes [13]. Some of the critical signaling pathways involved in HCC include, PI3K/AKT signaling, mammalian target of rapamycin (mTOR) signaling, Wnt/β-catenin, Ras/Raf/MAPK signaling, JAK/STAT signaling, Hippo signaling pathway governed by various receptor tyrosine kinase signaling modules (RTKs) [14–26]. These signaling pathways play an important role in governing various aspects of tumor cell behavior including differentiation, proliferation, invasion, metastasis and angiogenesis. However, the precise mechanism of development and progression of HCC remain unclear.

The rapid advancement of high-throughput microarray analysis has significantly enhanced the ability to comprehend fundamental mechanisms and prevalent genetic changes associated with cancer initiation and progression [27–31]. These alterations include genomic instability, single nucleotide polymorphisms, somatic mutations implicated in the initiation and progression of HCC [28].These genetic changes are intricately intertwined within the elaborate framework of various complex signaling pathways. In addition to these individual signaling pathways, there exist multiple layers of crosstalk interactions among these pathways regulating each other [32–34]. There is a growing emphasis on exploring novel therapeutic targets that specifically target the molecular pathways associated with tumorigenesis [35, 36]. In the present study, a molecular interaction map of HCC was constructed from literature review. The network was further analyzed to identify the influential genes and explore their roles in driving the progression of HCC.

## Methods

### Data curation and network assembly

To comprehensively elucidate the regulatory mechanisms governing the development and progression of HCC, an elaborate literature survey was conducted to identify the diverse array of regulators (Supplementary Table S3), their interactions and signaling pathways involved [37]. Subsequently, Cell Designer 4.4 was utilized to construct the disease map of HCC. Cell Designer 4.4 is known for its adeptness in representing biological pathways [38]. It follows a standardized depiction approach for biological components, interactions and supports the SMBL (Systems Biology Markup Language) format [39]. This enables the seamless exchange of disease networks across different software applications for various analyses. To ensure the use of precise and standardized terminology, HUGO nomenclature was obtained from Human Gene Nomenclature Committee (HGNC) (https://www.genenames.org) for all the regulators included in the disease map.

### Functional Enrichment Analysis

Functional enrichment analysis of all the regulators involved in the assembled HCC molecular map was conducted through Gene Ontology (GO) term analysis. This encompasses the aspects of Biological Processes (BP), Molecular Functions (MF) and Cellular Components (CC). Enrichr (https://maayanlab.cloud/Enrichr/), a comprehensive web tool for the gene set enrichment was used to perform batch functional annotation of all the regulators. The development of GO made it possible to analyze gene lists in the context of existing knowledge [40]. The incorporation of GO facilitated the thorough examination of gene functions. GO terms with a significance level of p < 0.05 were considered statistically significant across all three categories.

### Identification of Hub genes

The assembled network was further visualized and analyzed using Cytoscape [41]. CytoHubba [42], a Cytoscape plugin was utilized to identify the highly connected subnetwork of the hepatocellular carcinoma (HCC) molecular map assembled. Cytohubba is tailored to detect the most influential nodes involved in the development and progression of HCC by specifically employing Maximal Clique Centrality (MCC) algorithm.

### Survival analysis, Validation and Genetic Alteration analysis

Gene expression profiling interactive analysis (GEPIA 2.0) was used to validate the prognostic relevance of identified hub genes [43]. Overall survival analysis was performed using the LIHC datasets. This includes the data of the normal (50 liver) and tumor (369 HCC) samples from the TCGA and GTEx databases. All the parameters were set to default value, and the cut-off value was quartile. Overall survival was evaluated based on the Mantel-Cox test with 95% confidence interval and Cox proportional hazardous ratio. Further, cBioPortal (http://www.cbioportal.org/) an extensive online database that offers comprehensive analysis utilizing multidimensional cancer genomic data was employed to analyze and visualize genomic data. A total of 1231 samples from 7 studies excluding the intrahepatic cholangiocarcinoma and adenomas were used to compare the genetic alterations involved among the identified hub genes in liver cancer. Statistical relevance was determined based on a log p-value of less than 0.05, for identification of prognostically relevant genes.

The step-by-step workflow detailing the systematic construction of Hepatocellular Carcinoma (HCC) and analysis of the assembled disease map followed for this study is shown in **Figure 1**. This integrated workflow systematically guides the exploration of the underlying molecular mechanisms, intricate interactions and critical hubs governing the progression of HCC.

**Figure 1.**
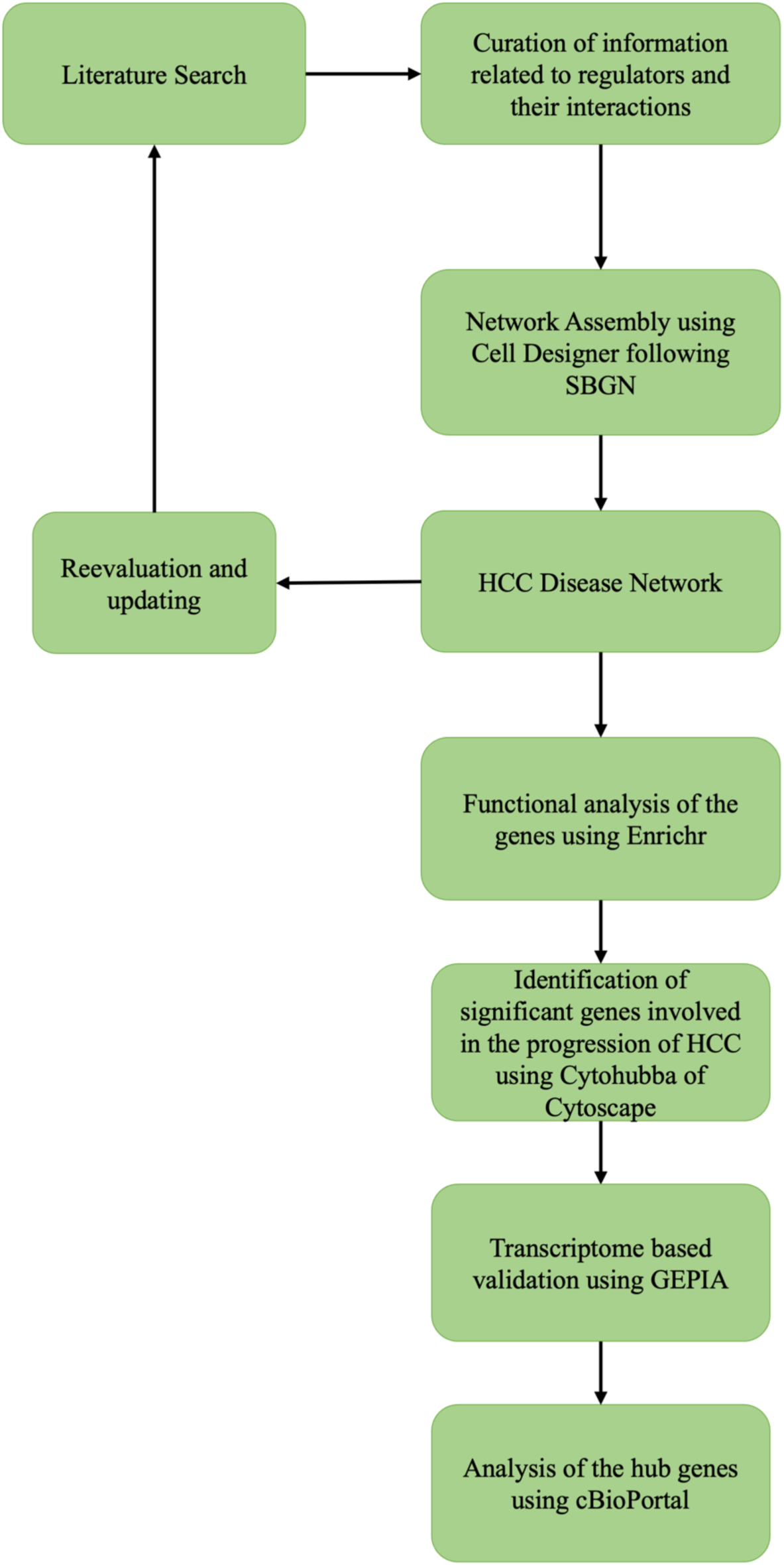
The workflow for the Construction of comprehensive map of Hepatocellular Carcinoma and analysis

## Results

### Molecular Interaction Map of HCC

A molecular map illustrating the regulators, their intricate signaling pathways, and underlying molecular mechanisms governing the development and progression of hepatocellular carcinoma (HCC) was assembled using Cell Designer V4.4 by conducting an extensive study of literature (**Figure 2**). The construction of this molecular map involved capturing cross-regulatory reactions spanning diverse signaling pathways. These include TGFβ signaling, FAK signaling, Jak/STAT signaling, Wnt/β-catenin signaling, GSK3β signaling, PI3K/Akt/mTOR signaling [14–26]. This assembled molecular map encompasses 193 distinct regulators (Supplementary Table S3) which includes a diverse range of molecular components including proteins, genes, RNA, and complexes. These components are interconnected through 362 reactions which outline the regulatory interactions. Among which 251 activating and 88 inhibiting reactions, 7 state transitions along with 15 complex associations were considered. These interactions were observed to give rise to three essential cellular states: proliferation, epithelial-mesenchymal transition (EMT), and apoptosis. The complete list of curated regulators, their HGNC, Uniprot, Ensembl IDs, along with the source of information are tabulated in **Supplementary File S2**. Additionally, the assembled HCC map strictly adheres to the Systems Biology Graphical Notation (SBGN) standards, ensuring the clear and standardized depiction of these complex regulatory relationships. Thus, this integrative approach allowed for the comprehensive depiction of molecular interactions that collectively contribute to the initiation, development, and progression of HCC.

**Figure 2.**
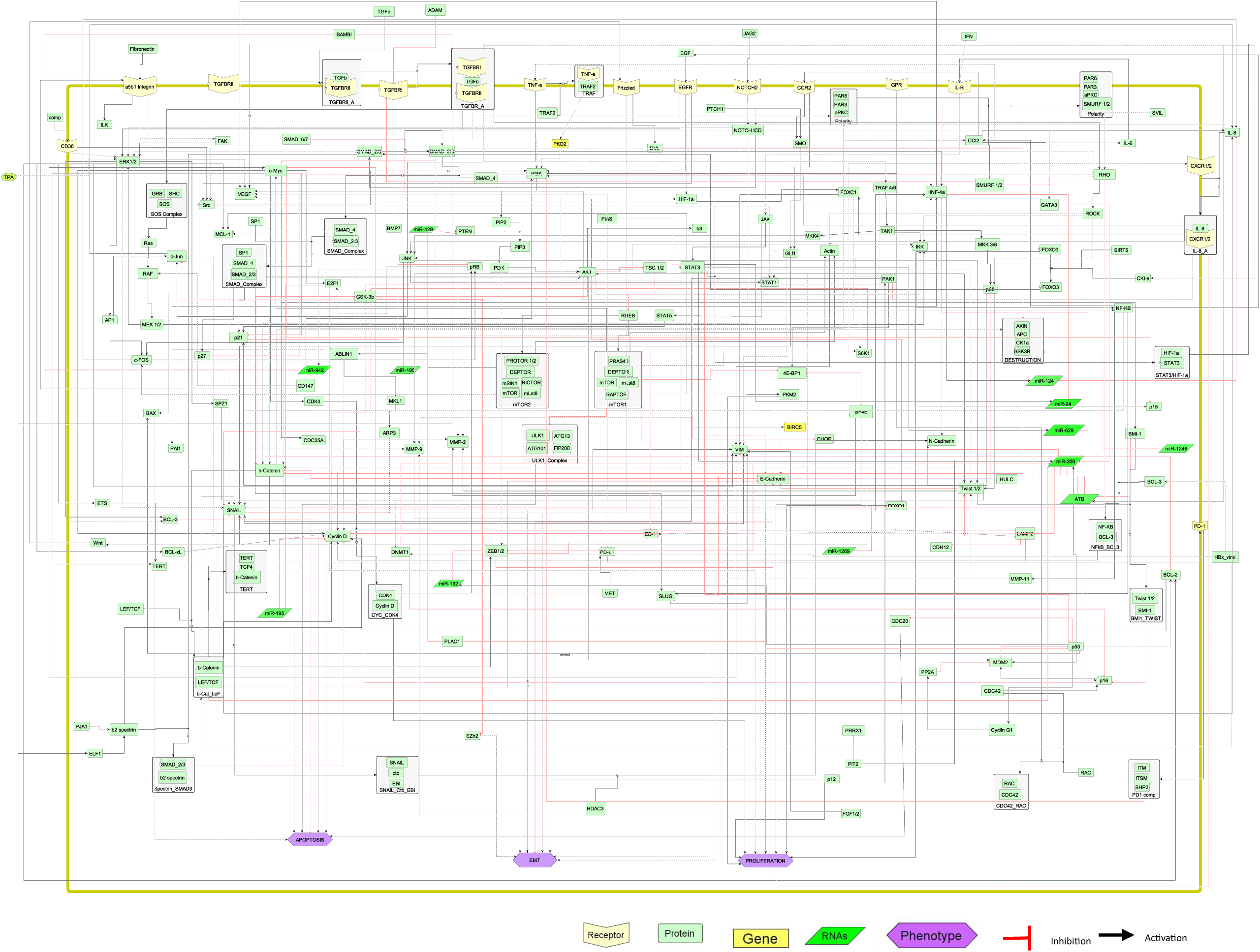
A molecular interaction map of HCC assembled using Cell Designer 4.4. The SBML compliant map assembled consists of 193 regulators and 362 interactions. The map illustrates the cross regulation between multiple signaling pathways including TGFβ, Wnt, Notch, JAK/STAT and Hypoxia signaling

### Functional Analysis of Identified Regulators

Enrichr online web tool was utilized to classify the gene ontology (GO) terms of the identified regulators from literature review (**Figure 3**). These regulators were primarily observed to be involved in various biological processes like regulation of cell proliferation, phosphorylation, cellular processes, regulation of apoptotic processes, migration, EMT (**Figure 3a**). Additionally, they were found to be associated with molecular functions like kinase activity and transcription factor binding, serine threonine activity, phosphatase binding (**Figure 3b**). Cellular component analysis revealed that these regulators were predominantly located at focal adhesion, cell substrate junctions, cytoskeleton, nucleus (**Figure 3c**). These findings indicate that the regulators play an active role in modulating a range of molecular functions, thereby influencing various signaling pathways and essential cellular processes, including cellular structures and proliferation.

**Figure 3.**
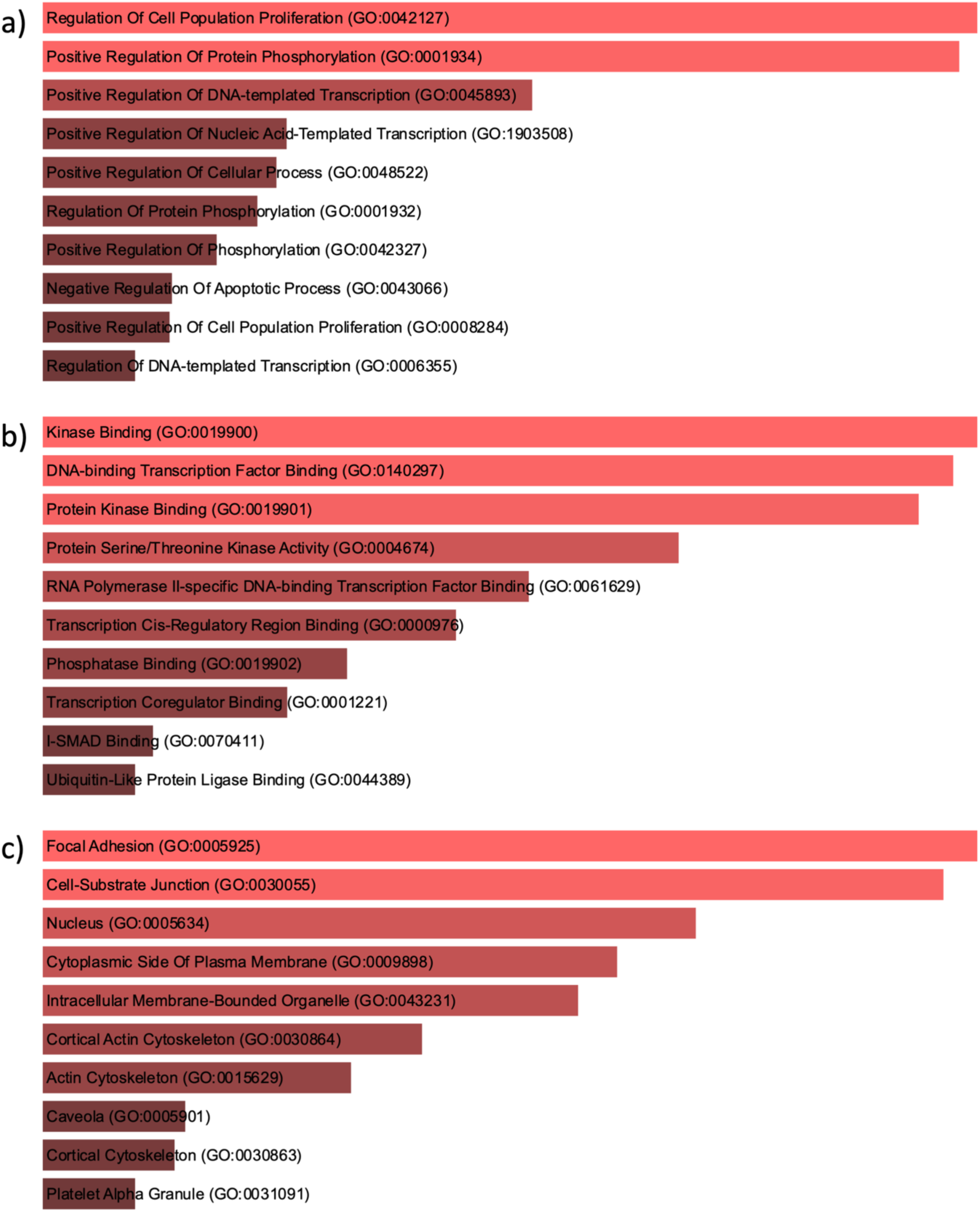
GO term enrichment analysis performed using Enrichr for the genes involved. The assembled maps top 10 (a) Biological processes (b) Molecular Functions, and (c) Cellular Components were represented.

Further, Venn webtool from bioinformatics and evolutionary genomics (https://bioinformatics.psb.ugent.be/webtools/Venn/) was used to identify the common genes involved in significant GO biological processes like Proliferation, EMT, apoptosis and migration (**Figure 4**). PTK2, IL6, CTNNB1, were observed to be common genes among proliferation, EMT, apoptosis while PTK2, AKT1, TJP1, VEGFA, AKT2 were observed to be the common genes among proliferation, migration, apoptosis. PTK2, emerges as a central player in both the sets of common genes, featuring its importance in orchestrating various aspects in the progression of HCC. Additionally, the identified common genes also indicate the delicate balance between cell growth, cell migration, and cell death in HCC.

**Figure 4.**
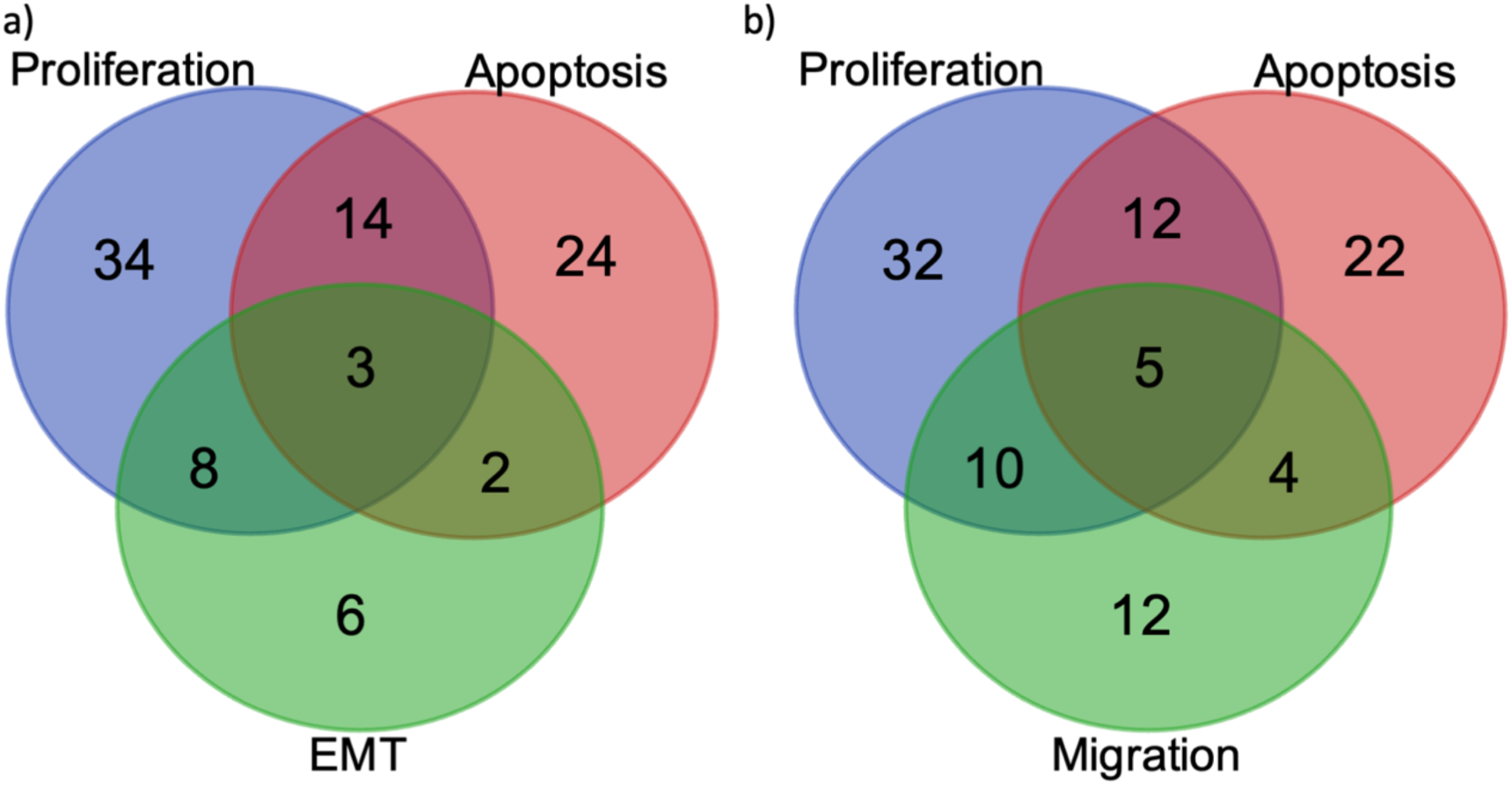
Commonly expressed genes among different biological processes. (a) PTK2, IL6, CTNNB1, were found to be commonly observed among proliferation, apoptosis and EMT, (b) PTK2, AKT1, TJP1, VEGFA, AKT2 were found to be commonly observed among proliferation, apoptosis and migration.

### Extraction of Subnetwork and identification of hub genes using Cytoscape: Cytohubba

Analysis of the network in Cytoscape using the CytoHubba plugin (**Figure 5**) led to the identification of the highly connected subnetwork (**Figure 6a**). This extracted network of top 20 nodes comprised one primary connected component and two sub-connected components. As depicted in the expanded subgraph (**Figure 6b**), these components were observed to be highly interconnected, influencing disease progression thorough crucial processes such as proliferation and EMT hence we considered them to be most influential regulators of the map (Supplementary Table S2). SNAIL1, ERK1, GSK3β, cMyc, STAT3, Twist1_2, CyclinD1, b_Catenin_LeF complex, miR200, VIM, AKT, E_cadherin, TGFBR_A Complex, JNK, HNF4a, p53, b_Catenin, and PI3K, were among the top 20 influential genes. The gradient from red to yellow implies the highly significant genes to significant genes. The presence of these diverse range of hub genes highlights their significance in orchestrating a multitude of cellular processes such as proliferation and EMT suggesting their involvement in the onset, development and progression of HCC.

**Figure 5.**
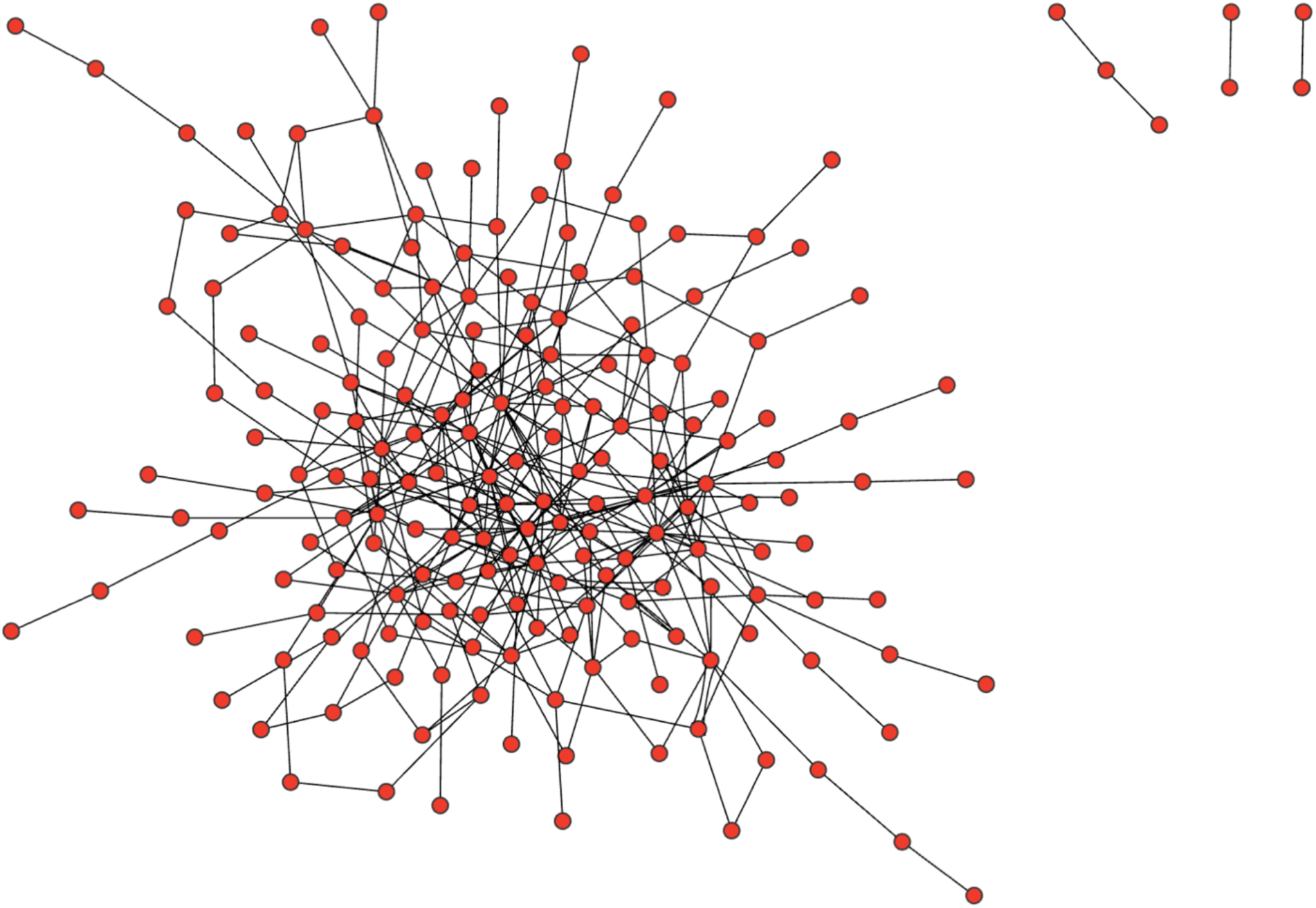
The network map of HCC visualized using the Compound Spring Embedder (CoSE) layout within the Cytoscape. This visualization highlights the intricate relationship of the components within the network. The network was visualized as a complex structure comprising one centrally connected component and 3 separate subgraphs.

**Figure 6.**
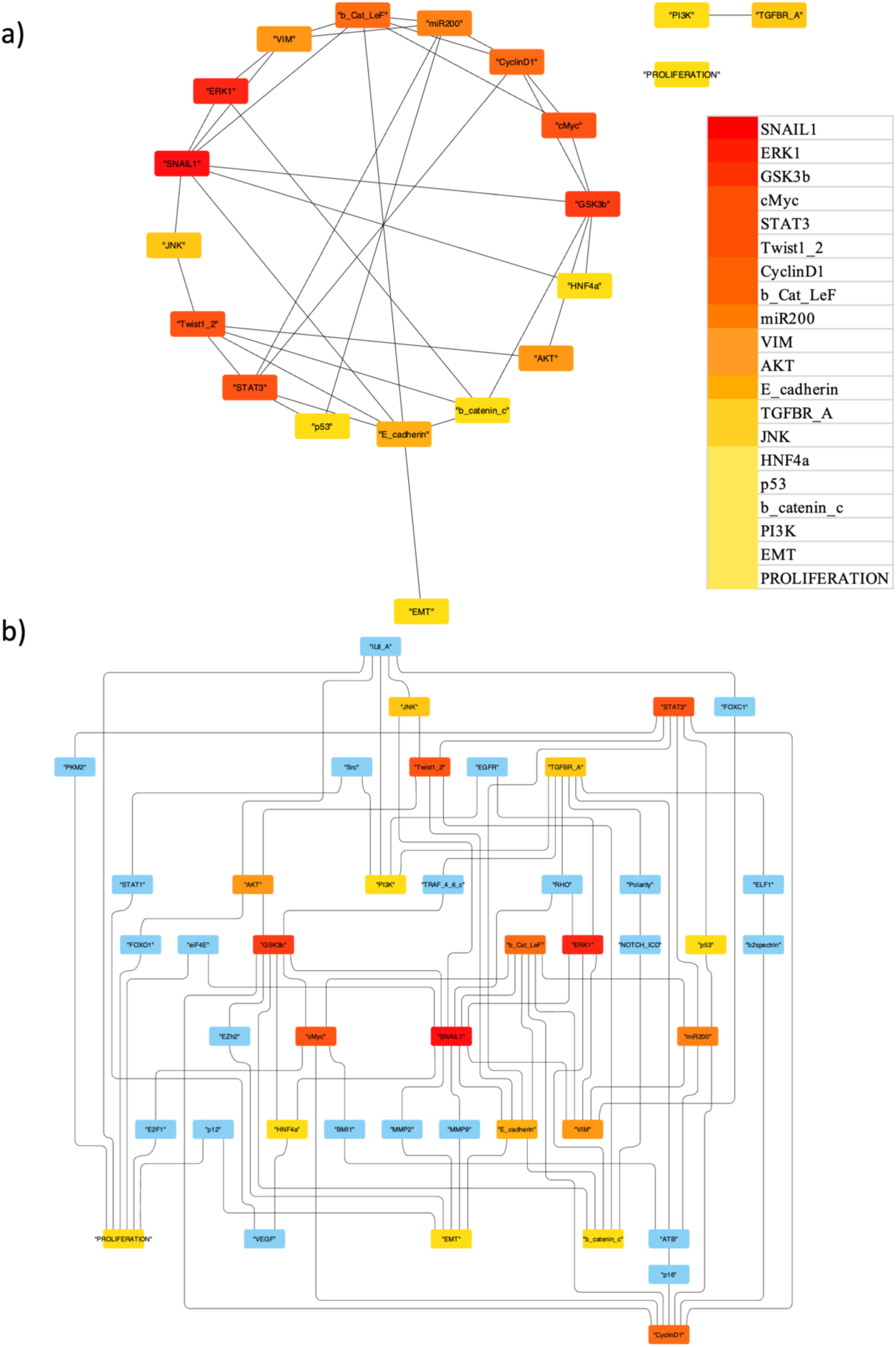
Identification of influential genes involved in HCC progression using cytohubba of Cytoscape (a)Top 20 significant hub genes were identified using MCC algorithm. Hub genes were defined as genes with the highest degree of connectivity within the network. The gradient from red to yellow implies the highly significant genes to significant genes (b) Expanded subnetwork illustrating the interconnected process identified through Cytohubba. This visualization illustrates the interconnectedness of the highly significant hub genes driving biological processes such as proliferation and EMT during HCC.

### Prognostic, genome analysis and validation of identified hub genes

The prognostic relevance of identified hub genes in HCC and their clinical relevance were further assessed using overall survival analysis. Liver Hepatocellular Cancer (LIHC) datasets from TCGA and GTEx project were evaluated using the GEPIA database (**Figure 7, Supplementary Figure S1**). PI3K, CTNNB1, AKT1, TGFB1, TWIST1, MAPK3, exhibited significant prognostic difference in overall survival with p<0.05. Additionally, as shown in **Figure 7** elevated expression of CTNNB1 was associated with better overall survival rates among LIHC patients compared to those with lower expressions. Further, genome alterations were examined using the TCGA cohort of HCC (N = 1231) using the cBioPortal database. The oncoprint results of the genome alterations of the identified hub genes were shown in **Figure 8a**. Notably, the transcription factor and tumor suppressor gene TP53 exhibited a higher frequency of alterations including driver mutations and deep deletions. Similarly, CTNNB1 showed frequencies for driver mutations and splice mutations. CCND1, Myc, PI3K, TGFβ displayed amplification frequencies in HCC datasets. AKT exhibited both amplifications (AKT2, AKT3) and deletions (AKT1), while GSK3β had frequencies for deep deletions. Conversely, the TWIST family of transcription factors displayed the least frequency of genome alterations (**Figure 8a)**. These observed genetic alterations were found to impact both the overall survival (p = 0.0137) and disease-free survival (p = 0.0213) of HCC patients (**Figure 8 b, c**). Specifically, patients belonging to unaltered genes group exhibited significant prognostic difference in survival to those in the altered genes group. These finding underscores the potential clinical significance of these genetic alterations which may influence the disease prognosis and patient outcomes in HCC.

**Figure 7.**
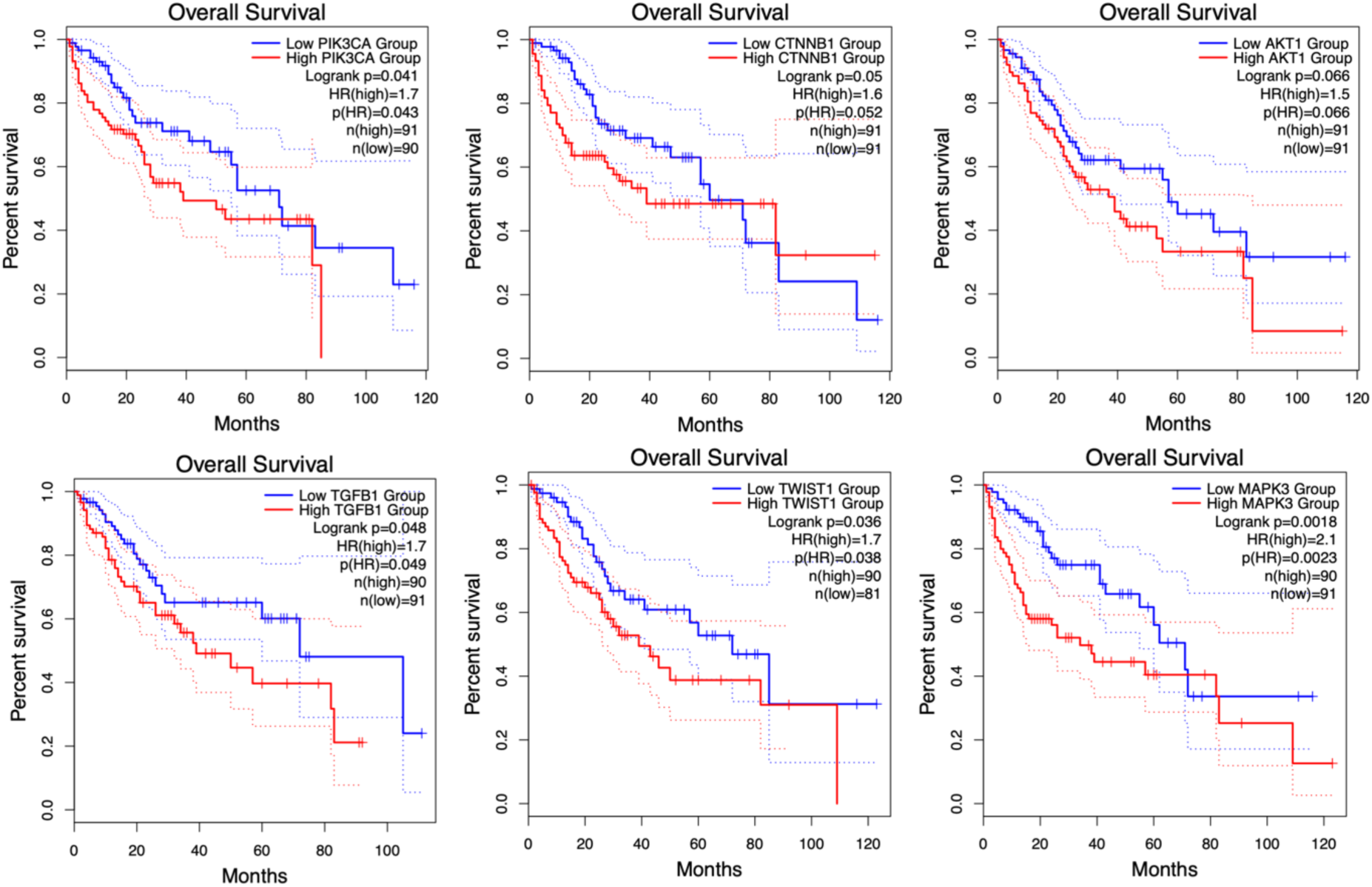
Prognostic significance of the identified hub genes in HCC patients. The solid lines of the plots represent the survival rate and dotted lines represent the 95% confidence interval of the same. A log p-value of less than 0.05 is considered prognostically significant. The overall conclusions from survival analysis evaluated using GEPIA indicates that genes PI3K, CTNNB1, AKT1, TGFB1, TWIST1, and MAPK3 were observed to be prognostically significant.

**Figure 8.**
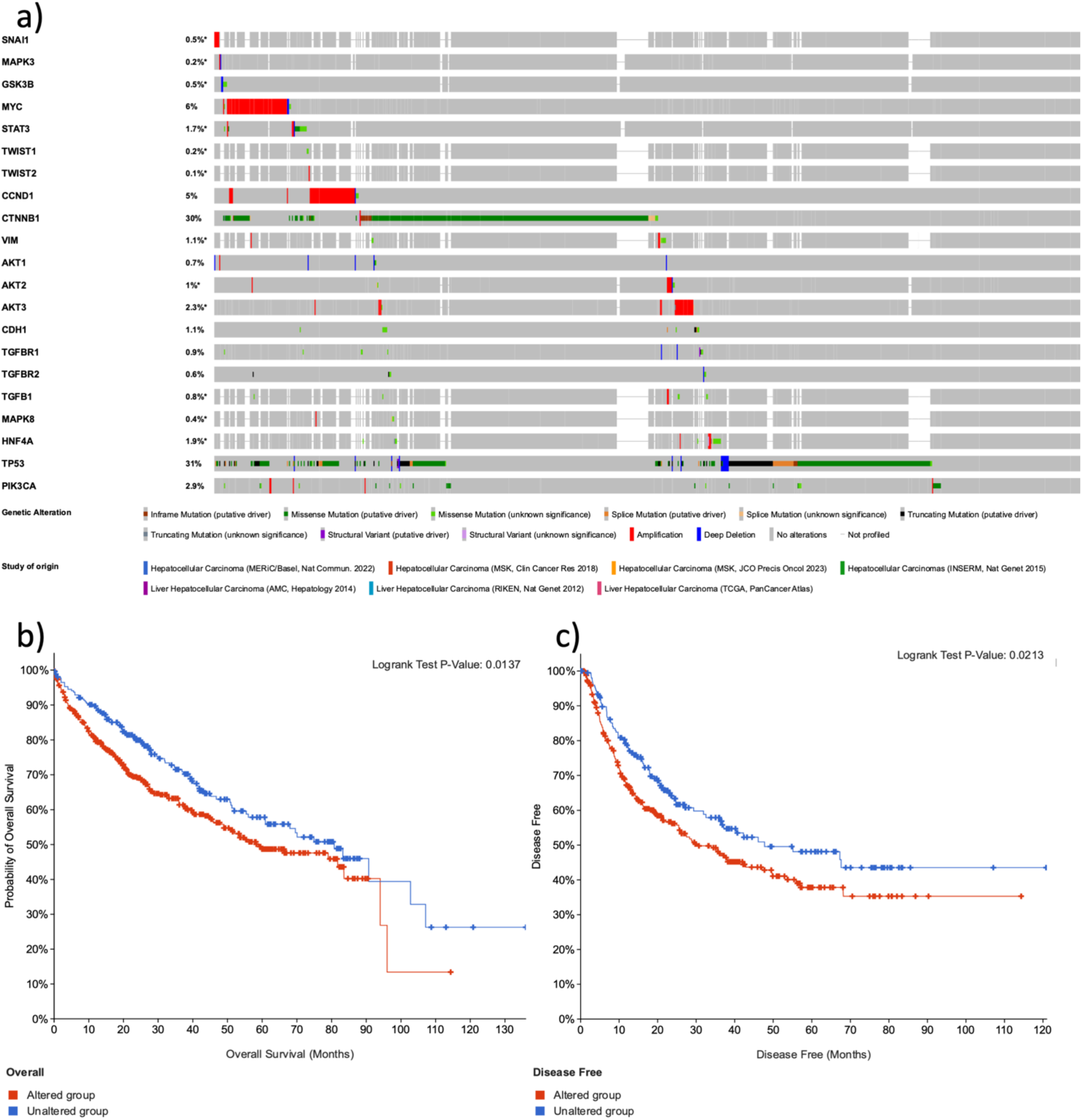
The genetic alteration analysis of the hub genes was identified using cBioPortal. (a) Oncoprint tab of the summary alterations in HCC datasets. TP53 was the gene with a significant number of alterations followed by CTNNB1 and CCND1; TWIST2 was the gene with less alterations, (b,c) Kaplan Meier plot of overall survival and Disease-free survival of the hub genes with and without gene alterations.

## Discussions

The current study explores the key regulators, signaling pathways and their molecular mechanisms implicated in the initiation, development and progression of Hepatocellular carcinoma (HCC). A molecular interaction map of HCC was assembled through an elaborate literature review. This assembled map depicts the intricate network of signaling pathways and regulators that participate in the dynamic continuum of HCC development and progression. The map captures several dysregulated signaling pathways including Wnt/β-catenin, TGFβ, JAK/STAT3, VEGF, crosstalk between them in HCC leading to uncontrolled cell growth and metastasis (**Figure 2**). Furthermore, these signaling pathways can result in the regulation of common target genes like FAK, CTNNB1, CCND1, GSK3β, PI3K, JNK which can regulate several biological processes and molecular functions contributing to the progression of HCC [44, 45]. This understanding of the molecular landscape of HCC is crucial for addressing the challenges in diagnosis and ultimately improving the treatment outcomes for individuals affected by this complex disease.

Molecular classification analysis of identified regulators using GO terms have disclosed that these regulators were primarily associated with the biological processes, functional processes that result in the genesis of HCC (**Figure 3a, b**). These regulators were localized to focal adhesion, cell substrate junction followed by nucleus maintaining tissue integrity and function (**Figure 3c**). Among these regulators, PTK2 was observed to be crucial for being common among various biological processes such as EMT, proliferation, apoptosis, and migration (**Figure 4**). PTK2 (FAK), a tyrosine kinase activated by several growth factors and integrins by phosphorylation targets multiple downstream signaling pathways in regulating different cellular functions [46, 47]. Several experimental evidence indicate that the over expression of PTK2 was observed in HCC. PTK2 was often over expressed concomitantly to β-catenin (CTNNB1) alterations in the context of HCC [48–51]. This further targets the downstream signaling pathways like PI3K signaling, AR signaling and many target genes like Myc, ERK in a PTK2/β-catenin dependent manner to regulate cell proliferation, survival, and migration in HCC [48, 49, 51].

Subsequent analysis involving extraction of subnetwork revealed broad spectrum of significant regulators in HCC (**Figure 6**). These hub genes were known to play a crucial role as transcription regulators, epithelial regulators, mesenchymal regulators, cell cycle regulators and regulators of cell proliferation (**Supplementary Table S2**). Among the identified genes PI3K, CTNNB1, AKT1, TGFB1, TWIST1, MAPK3 were found to be most significant in the overall survival analysis between the normal and cancer samples (p<0.05) (**Figure 7**).

GSK3β, a serine-threonine kinase was one of the highly significant hub genes identified from the disease map assembled (**Figure 6**). It was known for its role in phosphorylating various target proteins, leading to their inactivation through mechanisms such as degradation or subcellular relocalization. [52]. Functioning at the crossroads of many signaling pathways GSK3β, exhibits context dependent dual role as both tumor suppressor and tumor promotor in HCC [53–55] and one of the well explored pathway is PI3K/AKT/GSK3β/PTEN cascade [56–58]. GSK3β was known for its tumor suppressor role and was also known to inhibit CCND1, an important regulator for G1 to S phase transition in cell cycle [59–61]. GSK3β was also known to stabilize SNAIL1 by phosphorylation and its sub cellular localization further inhibiting EMT which is abrogated by hepatitis virus or by GSK3β inhibition [62, 63]. It directly inhibits FAK through phosphorylation, impeding cell migration. Interestingly, FAK was observed to evade this inhibition by phosphorylating GSK3β [55, 64] indicating complex interplay within these signaling networks.

On the contrary, GSK3β positively regulates NF-kB in cancer promoting HCC [65–69]. Furthermore β-catenin was also targeted by GSK3β in Wnt pathway in regulating cell adhesion and migration [70, 71]. Mutations in β-catenin bypasses the GSK3β degradation thus activating Wnt/β-catenin pathway leading to EMT and metastasis of HCC [72, 73]. Consequently, β-catenin regulates its direct target genes such as Myc, CCND1, ZEB1 in further promoting the uncontrolled progression of cell cycle [74–76]. It was speculated that GSK3β promotes metastasis by stabilizing PTEN and enhancing the phosphorylation of FAK, consequently accelerating the secretion of MMP-2 and MMP-9 [77]. Additionally, studies have also shown that inhibition of GSK3β decreased the FAK (PTK2) phosphorylation during cell migration and spread. Thus, the collective influence of various identified significant regulators and their signaling mechanisms impact over various cellular processes in the progression of HCC.

Analysis of genome alterations revealed varying frequencies of alterations among the identified hub genes. Among which TP53 and CTNNB1 emerged as highly altered genes (**Figure 8**). In contrast, no alterations were detected in TWIST2 (**Figure 8a**). The analysis of gene alterations unveiled a notable difference in overall survival and disease-free survival between the unaltered and altered groups (**Figure 8b**). Frequent occurrences of somatic mutations in TP53 and CTNNB1 were shown in previous reports [78–84]. The results of the present study corroborate with these findings, highlighting TP53 and CTNNB1 as the most frequently mutated genes in HCC [85]. These mutated driver genes were observed to be associated with cell cycle pathway and Wnt/β-catenin pathway [82, 84, 86]. Moreover, the association of these genetic alterations with overall patient survival underscores the complex interplay between genomic changes and disease progression in HCC. These mutations exert a substantial influence on the modulation of critical signaling pathways governing the development and progression of HCC. Further investigations and comprehensive understanding of the mechanistic implications of these alterations are warranted to elucidate their precise roles in HCC development and progression.

## Conclusions

HCC is a highly heterogenic cancer involving complex pathogenesis driven by various genetic factors and genomic alterations responsible for tumor proliferation and growth. This involves intricate signaling pathways, with multiple layers of interconnected interactions regulating each other in the progression of HCC. In the present study, a comprehensive map for the initiation, development and progression of HCC was developed and analyzed. Together results from the molecular functional analysis of gene ontology term biological processes, subgraph analysis using Cytoscape, prognosis analysis and genome alteration analysis clearly indicates that GSK3β, PTK2 (FAK), β-catenin are crucial players in the development and progression of HCC. Thus, the integrated workflow of assembling, analyzing the disease maps of HCC provides insights and comprehensive understanding of the disease mechanism. These findings highlight the prognostic and clinical significance of the various identified regulators in influencing the disease prognosis and patient outcomes in HCC.

## Supplementary Materials

The Supplementary File 1 contains the Supplementary Figure S1 and Supplementary Table S2

The Supplementary File 2 contains Supplementary Table S3

## Data Availability Statement

The assembled HCC map is available in the GitHub repository: https://github.com/gsb-sai/Hepatocellular-Carcinoma

## Author Contributions

AP and GSB designed the work; GSB and AM performed the modelling and analysis study; AP and GSB wrote the article.

## Funding

This research received no external funding.

## Acknowledgements

GSB, AM and AP acknowledge the facilities and support extended by NIT W, India.

## Conflicts of Interest

The authors declare no conflict of interest.

## Notes

### Competing Interest Statement

The authors have declared no competing interest.

