## Supplementary File S1 for "Comprehensive Network of Signaling Pathways in Hepatocellular Carcinoma and Network Analysis Reveals PTK2, GSK3β, and β-catenin to be Crucial for Cancer Progression"

### Supplementary File 1 for

### Comprehensive Network of Signaling Pathways in Hepatocellular Carcinoma and Network Analysis Reveals PTK2, GSK3β, and β-catenin to be Crucial for Cancer Progression

**Sai Bhavani Gottumukkala [1], Akilan M [2] and Anbumathi Palanisamy [3,*]**

[1] Research Scholar, Department of Biotechnology, National Institute of Technology Warangal, India;
[2] Post Graduate student, Department of Biotechnology, National Institute of Technology Warangal, India;
[3] Assistant Professor, Department of Biotechnology, National Institute of Technology Warangal, India

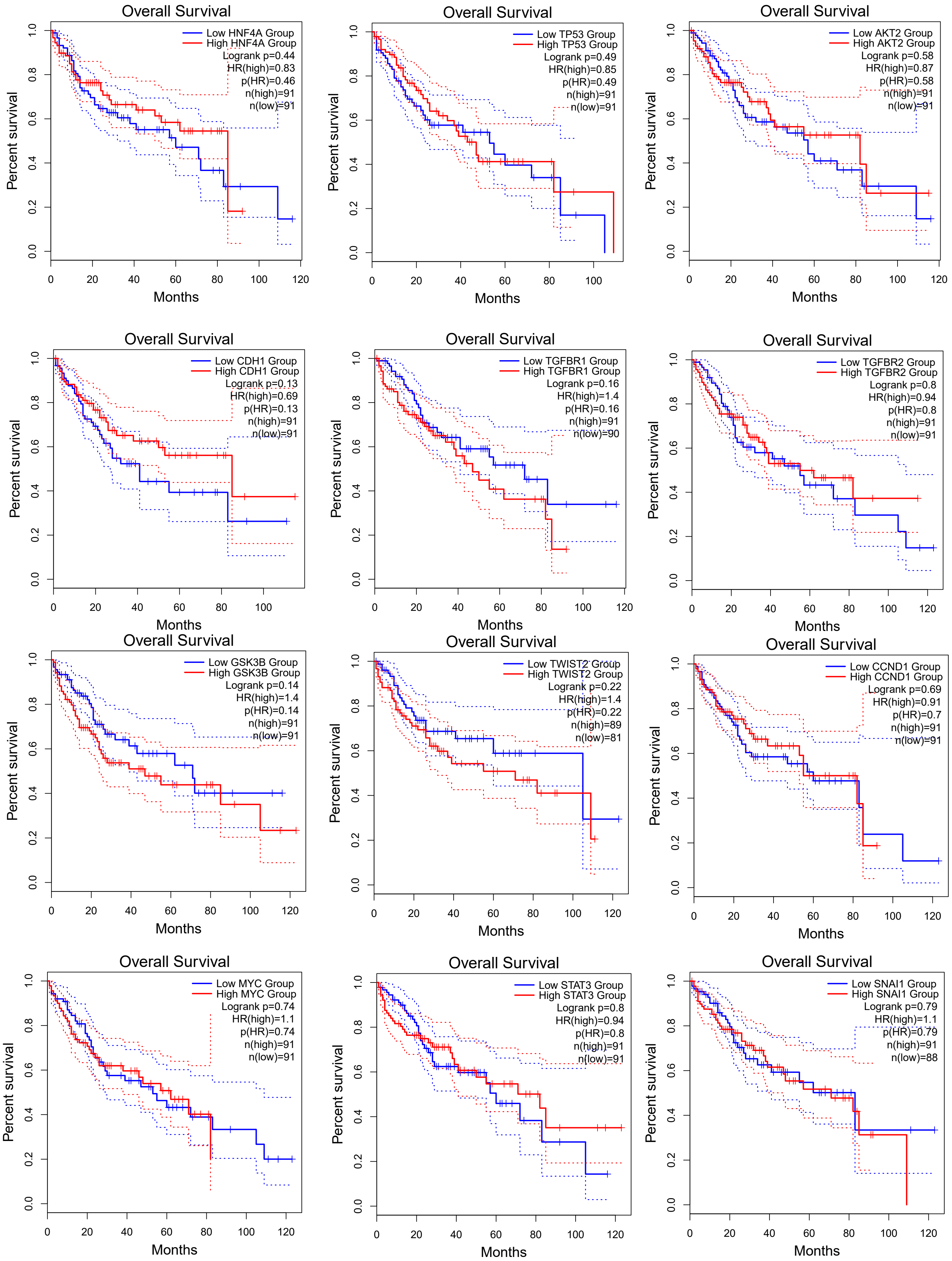

Overall Survival
Low HNF4A Group
High HNF4A Group
Logrank p=0.44
HR(high)=0.83
p(HR)=0.46
n(high)=91
n(low)=91
Percent survival
Months
Overall Survival
Low TP53 Group
High TP53 Group
Logrank p=0.49
HR(high)=0.85
p(HR)=0.49
n(high)=91
n(low)=91
Percent survival
Months
Overall Survival
Low AKT2 Group
High AKT2 Group
Logrank p=0.58
HR(high)=0.87
p(HR)=0.58
n(high)=91
n(low)=91
Percent survival
Months
Overall Survival
Low CDH1 Group
High CDH1 Group
Logrank p=0.13
HR(high)=0.69
p(HR)=0.13
n(high)=91
n(low)=91
Percent survival
Months
Overall Survival
Low TGFBR1 Group
High TGFBR1 Group
Logrank p=0.16
HR(high)=1.4
p(HR)=0.16
n(high)=91
n(low)=90
Percent survival
Months
Overall Survival
Low TGFBR2 Group
High TGFBR2 Group
Logrank p=0.8
HR(high)=0.94
p(HR)=0.8
n(high)=91
n(low)=91
Percent survival
Months
Overall Survival
Low GSK3B Group
High GSK3B Group
Logrank p=0.14
HR(high)=1.4
p(HR)=0.14
n(high)=91
n(low)=91
Percent survival
Months
Overall Survival
Low TWIST2 Group
High TWIST2 Group
Logrank p=0.22
HR(high)=1.4
p(HR)=0.22
n(high)=89
n(low)=81
Percent survival
Months
Overall Survival
Low CCND1 Group
High CCND1 Group
Logrank p=0.69
HR(high)=0.91
p(HR)=0.7
n(high)=91
n(low)=91
Percent survival
Months
Overall Survival
Low MYC Group
High MYC Group
Logrank p=0.74
HR(high)=1.1
p(HR)=0.74
n(high)=91
n(low)=91
Percent survival
Months
Overall Survival
Low STAT3 Group
High STAT3 Group
Logrank p=0.8
HR(high)=0.94
p(HR)=0.8
n(high)=91
n(low)=91
Percent survival
Months
Overall Survival
Low SNAI1 Group
High SNAI1 Group
Logrank p=0.79
HR(high)=1.1
p(HR)=0.79
n(high)=91
n(low)=88
Percent survival
Months

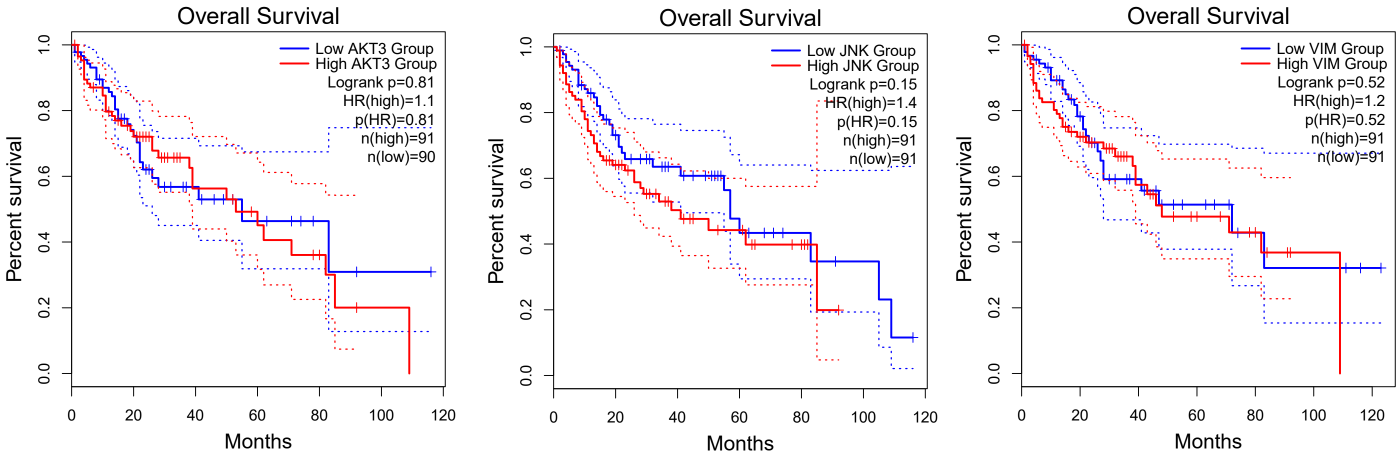


**Supplementary Figure S1**. Prognostic significance of the identified hub genes in HCC patients. The overall conclusions from survival analysis were evaluated using GEPIA for genes HNF4A, TP53, AKT2, CDH1, TGFBR1, TGFBR2, GSK3B, TWIST2, CCND1, MYC, STAT3, SNAI1, AKT3, JNK, VIM. A log p-value of less than 0.05 is considered to identify the prognostically significant genes.

**Table S2. specific roles of Hub regulators identified**

| **EMT Regulators** | **Oncogenic Signaling** | **Oncogenes** | **Tumor Suppressors** |
|---|---|---|---|
| SNAIL1 | PI3K | β-Catenin | p53 |
| TWIST | ERK | c-Myc | GSK3β |
| TGFβR | STAT3 | CyclinD1 | |
| VIM | JNK | AKT | |
| E-Cadherin | HNF4$\alpha$ | | |
| miR200 | LEF | | |
