## Supplementary File S2 for "Comprehensive Network of Signaling Pathways in Hepatocellular Carcinoma and Network Analysis Reveals PTK2, GSK3β, and β-catenin to be Crucial for Cancer Progression"

**Supplementary File 2 for**

**Comprehensive Network of Signaling Pathways in Hepatocellular Carcinoma and Network Analysis Reveals PTK2, GSK3β, and β-catenin to be Crucial for Cancer Progression**

**Sai Bhavani Gottumukkala [1], Akilan M [2] and Anbumathi Palanisamy [3,*]**
[1] Research Scholar, Department of Biotechnology, National Institute of Technology Warangal, India;
[2]Post Graduate student, Department of Biotechnology, National Institute of Technology Warangal, India;
[3] Assistant Professor, Department of Biotechnology, National Institute of Technology Warangal, India

**Supplementary Table S3.** List of regulators curated from the literature review that are involved in the development and progression of HCC

| Input | Approved symbol | Approved name | HGNC ID | Uniport ID/miRbase | Ensembl | RefSeq | References |
|---|---|---|---|---|---|---|---|
| BMI1 | BMI1 | BMI1 proto-oncogene, polycomb ring finger | HGNC:1066 | P35226 | ENSG00000168283 | NM_005180 | [1, 2] |
| TWIST | TWIST1 | twist family bHLH transcription factor 1 | HGNC:12428 | Q15672 | ENSG00000122691 | NM_000474 | [3-7] |
| | TWIST2 | twist family bHLH transcription factor 2 | HGNC:20670 | Q8WVJ9 | ENSG00000233608 | NM_001271893 | |
| CDC42 | CDC42 | cell division cycle 42 | HGNC:1736 | P60953 | ENSG00000070831 | NM_001791 | [8-10] |
| RAC | RAC1 | Rac family small GTPase 1 | HGNC:9801 | P63000 | ENSG00000136238 | NM_018890 | [11-13] |
| CCND1 | CCND1 | cyclin D1 | HGNC:1582 | P24385 | ENSG00000110092 | NM_053056 | [14-16] |

| | | | | | | | |
|---|---|---|---|---|---|---|---|
| CDK4 | CDK4 | cyclin dependent kinase 4 | HGNC:1773 | P11802 | ENSG0000013 5446 | NM_000075 | [16, 17] |
| AXIN | AXIN1 | axin 1 | HGNC:903 | O15169 | ENSG0000010 3126 | NM_003502 | [18, 19] |
| | AXIN2 | axin 2 | HGNC:904 | Q9Y2T1 | ENSG0000016 8646 | NM_004655 | |
| APC | APC | APC regulator of WNT signaling pathway | HGNC:583 | P25054 | ENSG0000013 4982 | NM_000038 | [20-22] |
| CK1a | CSNK1A1 | casein kinase 1 alpha 1 | HGNC:2451 | P48729 | ENSG0000011 3712 | NM_001892 | [23] |
| GSK3B | GSK3B | glycogen synthase kinase 3 beta | HGNC:4617 | P49841 | ENSG0000008 2701 | NM_00114615 6 | [23-25] |
| IL8 | CXCL8 | C-X-C motif chemokine ligand 8 | HGNC:6025 | P10145 | ENSG0000016 9429 | NM_000584 | [26-28] |
| CXCR | CXCR1 | C-X-C motif chemokine receptor 1 | HGNC:6026 | P25024 | ENSG0000016 3464 | NM_000634 | |
| | CXCR2 | C-X-C motif chemokine receptor 2 | HGNC:6027 | P25025 | ENSG0000018 0871 | NM_001557 | |
| NFKB1 | NFKB1 | nuclear factor kappa B subunit 1 | HGNC:7794 | P19838 | ENSG0000010 9320 | NM_003998 | [29-33] |
| NFKB2 | NFKB2 | nuclear factor kappa B subunit 2 | HGNC:7795 | Q00653 | ENSG0000007 7150 | NM_00107749 4 | |
| SHP2 | PTPN11 | protein tyrosine phosphatase non-receptor type 11 | HGNC:9644 | Q06124 | ENSG0000017 9295 | NM_00133043 7 | [34, 35] |
| PAR6 | PARD6A | par-6 family cell polarity regulator alpha | HGNC:15943 | Q9NPB6 | ENSG0000010 2981 | NM_016948 | [36-39] |
| PAR3 | PARD3 | par-3 family cell polarity regulator | HGNC:16051 | Q8TEW0 | ENSG0000014 8498 | NM_019619 | |

| | | | | | | | |
|---|---|---|---|---|---|---|---|
| PAR3 | F2RL2 | coagulation factor II thrombin receptor like 2 | HGNC:3539 | O00254 | ENSG00000164220 | NM_004101 | |
| aPKC | PRKCA | protein kinase C alpha | HGNC:9393 | P17252 | ENSG00000154229 | NM_002737 | [40, 41] |
| SMURF | SMURF1 | SMAD specific E3 ubiquitin protein ligase 1 | HGNC:16807 | Q9HCE7 | ENSG00000198742 | NM_020429 | [42, 43] |
| | SMURF2 | SMAD specific E3 ubiquitin protein ligase 2 | HGNC:16809 | Q9HAU4 | ENSG00000108854 | NM_022739 | |
| SMAD2 | SMAD2 | SMAD family member 2 | HGNC:6768 | Q15796 | ENSG00000175387 | NM_005901 | [27, 29, 44-52] |
| SMAD3 | SMAD3 | SMAD family member 3 | HGNC:6769 | P84022 | ENSG00000166949 | NM_005902 | |
| SMAD4 | SMAD4 | SMAD family member 4 | HGNC:6770 | Q13485 | ENSG00000141646 | NM_005359 | |
| SNAIL | SNAI1 | snail family transcriptional repressor 1 | HGNC:11128 | O95863 | ENSG00000124216 | NM_005985 | [23, 31, 53-56] |
| CTB | CTBP1 | C-terminal binding protein 1 | HGNC:2494 | Q13363 | ENSG00000159692 | NM_001328 | [57] |
| GRB | GRB2 | growth factor receptor bound protein 2 | HGNC:4566 | P62993 | ENSG00000177885 | NM_002086 | [58, 59] |
| SHCA | SHC1 | SHC adaptor protein 1 | HGNC:10840 | P29353 | ENSG00000160691 | NM_183001 | |
| SOS | SOS1 | SOS Ras/Rac guanine nucleotide exchange factor 1 | HGNC:11187 | Q07889 | ENSG00000115904 | NM_005633 | |
| STAT3 | STAT3 | signal transducer and activator of transcription 3 | HGNC:11364 | P40763 | ENSG00000168610 | NM_139276 | [60-64] |

| | | | | | | | |
|---|---|---|---|---|---|---|---|
| HIF1a | HIF1A | hypoxia inducible factor 1 subunit alpha | HGNC:4910 | Q16665 | ENSG00000100644 | NM_001530 | [65-67] |
| b2Spectrin | SPTB | spectrin beta, erythrocytic | HGNC:11274 | P11277 | ENSG00000070182 | NM_001024858 | [68] |
| TCF4 | TCF4 | transcription factor 4 | HGNC:11634 | P15884 | ENSG00000196628 | NM_003199 | [69] |
| CTNNB1 | CTNNB1 | catenin beta 1 | HGNC:2514 | P35222 | ENSG00000168036 | NM_001098210 | [15, 23, 24, 70-74] |
| TERT | TERT | telomerase reverse transcriptase | HGNC:11730 | O14746 | ENSG00000164362 | NM_001193376 | [75, 76] |
| TGFB | TGFB1 | transforming growth factor beta 1 | HGNC:11766 | P01137 | ENSG00000105329 | NM_000660 | [11, 17, 24, 44, 47, 48, 50, 61, 77-80] |
| | TGFB2 | transforming growth factor beta 2 | HGNC:11768 | P61812 | ENSG00000092969 | NM_003238 | |
| | TGFB3 | transforming growth factor beta 3 | HGNC:11769 | P10600 | ENSG00000119699 | NM_003239 | |
| BIRC5 | BIRC5 | baculoviral IAP repeat containing 5 | HGNC:593 | O15392 | ENSG00000089685 | NM_001168 | [81] |
| PKD2 | PRKD2 | protein kinase D2 | HGNC:17293 | Q9BZL6 | ENSG00000105287 | NM_016457 | [82] |
| 4E-BP1 | EIF4EBP1 | eukaryotic translation initiation factor 4E binding protein 1 | HGNC:3288 | Q13541 | ENSG00000187840 | NM_004095 | [83, 84] |
| ADAM | ADAM17 | ADAM metallopeptidase domain 17 | HGNC:195 | P78536 | ENSG00000151694 | NM_001382777 | [85] |
| AKT1 | AKT1 | AKT serine/threonine kinase 1 | HGNC:391 | P31749 | ENSG00000142208 | NM_005163 | [11, 17, 24, 72, 86] |

| | | | | | | | |
|---|---|---|---|---|---|---|---|
| AKT2 | AKT2 | AKT serine/threonine kinase 2 | HGNC:392 | P31751 | ENSG00000105221 | NM_001626 | |
| AKT3 | AKT3 | AKT serine/threonine kinase 3 | HGNC:393 | Q9Y243 | ENSG00000117020 | NM_181690 | |
| AP1 | | | | | | | [17, 87, 88] |
| ARP3 | ACTR3 | actin related protein 3 | HGNC:170 | P61158 | ENSG00000115091 | NM_005721 | [89, 90] |
| Actin | ACTB | actin beta | HGNC:132 | P60709 | ENSG00000075624 | NM_001101 | [90] |
| BAMBI | BAMBI | BMP and activin membrane bound inhibitor | HGNC:30251 | Q13145 | ENSG00000095739 | NM_012342 | [91, 92] |
| BAX | BAX | BCL2 associated X, apoptosis regulator | HGNC:959 | Q07812 | ENSG00000087088 | NM_138763 | [93] |
| BCL-2 | BCL2 | BCL2 apoptosis regulator | HGNC:990 | P10415 | ENSG00000171791 | NM_000633 | [54, 69, 93, 94] |
| BCL-3 | BCL3 | BCL3 transcription coactivator | HGNC:998 | P20749 | ENSG00000069399 | NM_005178 | |
| BCL-xL | BCL2L1 | BCL2 like 1 | HGNC:992 | Q07817 | ENSG00000171552 | NM_138578 | [54, 69] |
| BMP7 | BMP7 | bone morphogenetic protein 7 | HGNC:1074 | P18075 | ENSG00000101144 | NM_001719 | [95, 96] |
| CCR2 | CCR2 | C-C motif chemokine receptor 2 | HGNC:1603 | P41597 | ENSG00000121807 | NM_000647 | [82, 97, 98] |
| CCl2 | CCL2 | C-C motif chemokine ligand 2 | HGNC:10618 | P13500 | ENSG00000108691 | NM_002982 | |
| CD147 | BSG | basigin (Ok blood group) | HGNC:1116 | P35613 | ENSG00000172270 | NM_001728 | [99-101] |

| | | | | | | | |
|---|---|---|---|---|---|---|---|
| CD36 | CD36 | CD36 molecule (CD36 blood group) | HGNC:1663 | P16671 | ENSG00000135218 | NM_001001547 | [102, 103] |
| CDC20 | CDC20 | cell division cycle 20 | HGNC:1723 | Q12834 | ENSG00000117399 | NM_001255 | [104] |
| CDC25A | CDC25A | cell division cycle 25A | HGNC:1725 | P30304 | ENSG00000164045 | NM_001789 | [105] |
| CHOP | DDIT3 | DNA damage inducible transcript 3 | HGNC:2726 | P0DPQ6 | ENSG00000175197 | NM_004083 | [106, 107] |
| CKIe | CSNK1E | casein kinase 1 epsilon | HGNC:2453 | P49674 | ENSG00000213923 | NM_001894 | [23, 108] |
| Cyclin G1 | CCNG1 | cyclin G1 | HGNC:1592 | P51959 | ENSG00000113328 | NM_004060 | [77] |
| DNMT1 | DNMT1 | DNA methyltransferase 1 | HGNC:2976 | P26358 | ENSG00000130816 | NM_001379 | [16, 109, 110] |
| DVL1 | DVL1 | dishevelled segment polarity protein 1 | HGNC:3084 | O14640 | ENSG00000107404 | NM_004421 | [111] |
| CDH1 | CDH1 | cadherin 1 | HGNC:1748 | P12830 | ENSG00000039068 | NM_004360 | [109, 112-118] |
| E2F1 | E2F1 | E2F transcription factor 1 | HGNC:3113 | Q01094 | ENSG00000101412 | NM_005225 | [16] |
| EGF | EGF | epidermal growth factor | HGNC:3229 | P01133 | ENSG00000138798 | NM_001178130 | [33] |
| EGFR | EGFR | epidermal growth factor receptor | HGNC:3236 | P00533 | ENSG00000146648 | NM_005228 | |
| ELF1 | ELOF1 | elongation factor 1 | HGNC:28691 | P60002 | ENSG00000130165 | NM_032377 | [119] |
| ERK1 | MAPK3 | mitogen-activated protein kinase 3 | HGNC:6877 | P27361 | ENSG00000102882 | NM_001040056 | [3, 14, 23, 87, 120, 121] |

| | | | | | | | |
|---|---|---|---|---|---|---|---|
| ERK2 | MAPK1 | mitogen-activated protein kinase 1 | HGNC:6871 | P28482 | ENSG00000100030 | NM_002745 | |
| ETS1 | ETS1 | ETS proto-oncogene 1, transcription factor | HGNC:3488 | P14921 | ENSG00000134954 | NM_005238 | [56, 57] |
| ETS2 | ETS2 | ETS proto-oncogene 2, transcription factor | HGNC:3489 | P15036 | ENSG00000157557 | NM_001256295 | |
| EZh2 | EZH2 | enhancer of zeste 2 polycomb repressive complex 2 subunit | HGNC:3527 | Q15910 | ENSG00000106462 | NM_004456 | [122, 123] |
| FAK | PTK2 | protein tyrosine kinase 2 | HGNC:9611 | Q05397 | ENSG00000169398 | NM_005607 | [17, 124] |
| FGF1 | FGF1 | fibroblast growth factor 1 | HGNC:3665 | P05230 | ENSG00000113578 | NM_000800 | [125-127] |
| FGF2 | FGF2 | fibroblast growth factor 2 | HGNC:3676 | P09038 | ENSG00000138685 | NM_002006 | |
| FOXC1 | FOXC1 | forkhead box C1 | HGNC:3800 | Q12948 | ENSG00000054598 | NM_001453 | [118, 128, 129] |
| FOXO1 | FOXO1 | forkhead box O1 | HGNC:3819 | Q12778 | ENSG00000150907 | NM_002015 | |
| FOXO3 | FOXO3 | forkhead box O3 | HGNC:3821 | O43524 | ENSG00000118689 | NM_001415139 | |
| Fibronectin | FN1 | fibronectin 1 | HGNC:3778 | P02751 | ENSG00000115414 | NM_212476 | [48, 100, 126, 130-132] |
| Frizzled | FZD1 | frizzled class receptor 1 | HGNC:4038 | Q9UP38 | ENSG00000157240 | NM_003505 | [20, 133-135] |
| GATA3 | GATA3 | GATA binding protein 3 | HGNC:4172 | P23771 | ENSG00000107485 | NM_001002295 | [101, 136] |

| | | | | | | | |
|---|---|---|---|---|---|---|---|
| GLI1 | GLI1 | GLI family zinc finger 1 | HGNC:4317 | P08151 | ENSG00000111087 | NM_005269 | [137-140] |
| GPR | CMKLR2 | chemerin chemokine-like receptor 2 | HGNC:4463 | P46091 | ENSG00000183671 | NM_001098199 | [12, 141, 142] |
| HDAC3 | HDAC3 | histone deacetylase 3 | HGNC:4854 | O15379 | ENSG00000171720 | NM_003883 | [109] |
| HNF-4a | HNF4A | hepatocyte nuclear factor 4 alpha | HGNC:5024 | P41235 | ENSG00000101076 | NM_000457 | [143, 144] |
| HULC | HULC | hepatocellular carcinoma up-regulated long non-coding RNA | HGNC:34232 | | ENSG00000285219 | NR_004855 | [145] |
| IFN | IFNA1 | interferon alpha 1 | HGNC:5417 | P01562 | ENSG00000197919 | NM_024013 | [130, 146, 147] |
| IKK | IKBKB | inhibitor of nuclear factor kappa B kinase subunit beta | HGNC:5960 | O14920 | ENSG00000104365 | NM_001190720 | [147-149] |
| IL-6 | IL6 | interleukin 6 | HGNC:6018 | P05231 | ENSG00000136244 | NM_000600 | [62, 64, 68] |
| IL-8 | CXCL8 | C-X-C motif chemokine ligand 8 | HGNC:6025 | P10145 | ENSG00000169429 | NM_000584 | [26] |
| IL-6R | IL6R | interleukin 6 receptor | HGNC:6019 | P08887 | ENSG00000160712 | NM_000565 | [64, 68] |
| ILK | ILK | integrin linked kinase | HGNC:6040 | Q13418 | ENSG00000166333 | NM_004517 | [150] |
| JAG2 | JAG2 | jagged canonical Notch ligand 2 | HGNC:6189 | Q9Y219 | ENSG00000184916 | NM_002226 | [85, 151] |
| JAK | JAK1 | Janus kinase 1 | HGNC:6190 | P23458 | ENSG00000162434 | NM_002227 | [26, 62, 63, 152] |

| | | | | | | | |
|---|---|---|---|---|---|---|---|
| | JAK2 | Janus kinase 2 | HGNC:6192 | O60674 | ENSG00000096968 | NM_001322194 | |
| | JAK3 | Janus kinase 3 | HGNC:6193 | P52333 | ENSG00000105639 | NM_000215 | |
| JNK | MAPK8 | mitogen-activated protein kinase 8 | HGNC:6881 | P45983 | ENSG0000010764 | NM_001278547 | [30, 49, 153, 154] |
| LAMP2 | LAMP2 | lysosomal associated membrane protein 2 | HGNC:6501 | P13473 | ENSG00000005893 | NM_001122606 | [55] |
| LEF/TCF | LEF1 | lymphoid enhancer binding factor 1 | HGNC:6551 | Q9UJU2 | ENSG00000138795 | NM_001130713 | [134, 155] |
| MCL-1 | MCL1 | MCL1 apoptosis regulator, BCL2 family member | HGNC:6943 | Q07820 | ENSG00000143384 | NM_021960 | [156, 157] |
| MDM2 | MDM2 | MDM2 proto-oncogene | HGNC:6973 | Q00987 | ENSG00000135679 | NM_002392 | [158-160] |
| MEK | MAP2K3 | mitogen-activated protein kinase kinase 3 | HGNC:6843 | P46734 | ENSG00000034152 | NM_145109 | [87] |
| | MAP2K4 | mitogen-activated protein kinase kinase 4 | HGNC:6844 | P45985 | ENSG00000065559 | NM_00128143 | |
| MET | MET | MET proto-oncogene, receptor tyrosine kinase | HGNC:7029 | P08581 | ENSG00000105976 | NM_000245 | [53] |
| MKK4 | MAP2K4 | mitogen-activated protein kinase kinase 4 | HGNC:6844 | P45985 | ENSG00000065559 | NM_001281435 | [161] |
| MKL1 | MRTFA | myocardin related transcription factor A | HGNC:14334 | Q969V6 | ENSG00000196588 | NM_020831 | [162] |
| MMP-11 | MMP11 | matrix metallopeptidase 11 | HGNC:7157 | P24347 | ENSG00000099953 | NM_005940 | [11, 87, 139, 163, 164] |
| MMP-2 | MMP2 | matrix metallopeptidase 2 | HGNC:7166 | P08253 | ENSG00000087245 | NM_001127891 | |

| | | | | | | | |
|---|---|---|---|---|---|---|---|
| MMP-9 | MMP9 | matrix metallopeptidase 9 | HGNC:7176 | P14780 | ENSG00000100985 | NM_004994 | |
| N-Cadherin | CDH2 | cadherin 2 | HGNC:1759 | P19022 | ENSG00000170558 | NM_001792 | [25, 123, 130, 165-167] |
| NOTCH1 | NOTCH1 | notch receptor 1 | HGNC:7881 | P46531 | ENSG00000148400 | NM_017617 | [64, 85, 168, 169] |
| NOTCH2 | NOTCH2 | notch receptor 2 | HGNC:7882 | Q04721 | ENSG00000134250 | NM_024408 | |
| PAI1 | SERPINE1 | serpin family E member 1 | HGNC:8583 | P05121 | ENSG00000106366 | NM_000602 | [62, 170] |
| PAK1 | PAK1 | p21 (RAC1) activated kinase 1 | HGNC:8590 | Q13153 | ENSG00000149269 | NM_002576 | [171] |
| PD-1 | PDCD1 | programmed cell death 1 | HGNC:8760 | Q15116 | ENSG00000188389 | NM_005018 | [172-174] |
| PD-L1 | CD274 | CD274 molecule | HGNC:17635 | Q9NZQ7 | ENSG00000120217 | NM_014143 | |
| PDK | PDPK1 | 3-phosphoinositide dependent protein kinase 1 | HGNC:8816 | O15530 | ENSG00000140992 | NM_001261816 | [40, 175, 176] |
| PI3K | PIK3CA | phosphatidylinositol-4,5-bisphosphate 3-kinase catalytic subunit alpha | HGNC:8975 | P42336 | ENSG00000121879 | NM_006218 | [11, 72, 124, 177, 178] |
| PIAS | PIAS1 | protein inhibitor of activated STAT 1 | HGNC:2752 | O75925 | ENSG00000033800 | NM_001320687 | [179] |
| PIP2 | PSTPIP2 | proline-serine-threonine phosphatase interacting protein 2 | HGNC:9581 | Q9H939 | ENSG00000152229 | NM_024430 | [180-182] |
| PJA1 | PJA1 | praja ring finger ubiquitin ligase 1 | HGNC:16648 | Q8NG27 | ENSG00000181191 | NM_145119 | [83, 183] |

| | | | | | | | |
|---|---|---|---|---|---|---|---|
| PKM2 | PKM | pyruvate kinase M1/2 | HGNC:9021 | P14618 | ENSG00000067225 | NM_001206796 | [73, 184] |
| PLAC1 | PLAC1 | placenta enriched 1 | HGNC:9044 | Q9HBJ0 | ENSG00000170965 | NM_021796 | [185, 186] |
| PP2A | PPP2CA | protein phosphatase 2 catalytic subunit alpha | HGNC:9299 | P67775 | ENSG00000113575 | NM_002715 | [187, 188] |
| PRRX1 | PRRX1 | paired related homeobox 1 | HGNC:9142 | P54821 | ENSG00000116132 | NM_006902 | [189, 190] |
| PTCH1 | PTCH1 | patched 1 | HGNC:9585 | Q13635 | ENSG00000185920 | NM_000264 | [139, 191, 192] |
| PTEN | PTEN | phosphatase and tensin homolog | HGNC:9588 | P60484 | ENSG00000171862 | NM_000314 | [193-196] |
| RAF | RAF1 | Raf-1 proto-oncogene, serine/threonine kinase | HGNC:9829 | P04049 | ENSG00000132155 | NM_002880 | [93, 177, 197] |
| RHEB | RHEB | Ras homolog, mTORC1 binding | HGNC:10011 | Q15382 | ENSG00000106615 | NM_005614 | [198] |
| RHO | RHOD | ras homolog family member D | HGNC:670 | O00212 | ENSG00000173156 | NM_014578 | [199-201] |
| ROCK | ROCK1 | Rho associated coiled-coil containing protein kinase 1 | HGNC:10251 | Q13464 | ENSG00000067900 | NM_005406 | |
| Ras | KRAS | KRAS proto-oncogene, GTPase | HGNC:6407 | P01116 | ENSG00000133703 | NM_033360 | [177, 202, 203] |
| | HRAS | HRas proto-oncogene, GTPase | HGNC:5173 | P31249 | ENSG00000128652 | NM_006898 | |
| S6K1 | RPS6KB1 | ribosomal protein S6 kinase B1 | HGNC:10436 | P23443 | ENSG00000108443 | NM_003161 | [204] |
| SIRT6 | SIRT6 | sirtuin 6 | HGNC:14934 | Q8N6T7 | ENSG00000077463 | NM_001193285 | [118, 205, 206] |

| | | | | | | | |
|---|---|---|---|---|---|---|---|
| SLUG | SNAI2 | snail family transcriptional repressor 2 | HGNC:11094 | O43623 | ENSG00000019549 | NM_003068 | [87, 153, 189, 207, 208] |
| SMO | SMO | smoothened, frizzled class receptor | HGNC:11119 | Q99835 | ENSG00000128602 | NM_005631 | [137, 163, 209, 210] |
| SP1 | SP1 | Sp1 transcription factor | HGNC:11205 | P08047 | ENSG00000185591 | NM_001251825 | [46, 211-214] |
| SPZ1 | SPZ1 | spermatogenic leucine zipper 1 | HGNC:30721 | Q9BXG8 | ENSG00000164299 | NM_032567 | [215] |
| STAT1 | STAT1 | signal transducer and activator of transcription 1 | HGNC:11362 | P42224 | ENSG00000115415 | NM_007315 | [216, 217] |
| STAT5 | STAT5A | signal transducer and activator of transcription 5A | HGNC:11366 | P42229 | ENSG00000126561 | NM_003152 | |
| SVIL | SVIL | supervillin | HGNC:11480 | O95425 | ENSG00000197321 | NM_001323599 | [218, 219] |
| Src | SRC | SRC proto-oncogene, non-receptor tyrosine kinase | HGNC:11283 | P12931 | ENSG00000197122 | NM_005417 | [17, 220] |
| TAK1 | MAP3K7 | mitogen-activated protein kinase kinase kinase 7 | HGNC:6859 | O43318 | ENSG00000135341 | NM_145331 | [4, 68, 221, 222] |
| SMAD6 | SMAD6 | SMAD family member 6 | HGNC:6772 | O43541 | ENSG00000137834 | NM_005585 | [18, 27, 51] |
| SMAD7 | SMAD7 | SMAD family member 7 | HGNC:6773 | O15105 | ENSG00000101665 | NM_005904 | |
| TGFBR1 | TGFBR1 | transforming growth factor beta receptor 1 | HGNC:11772 | P36897 | ENSG00000106799 | NM_001130916 | [44, 45, 49, 51, 223, 224] |
| TGFBR2 | TGFBR2 | transforming growth factor beta receptor 2 | HGNC:11773 | P37173 | ENSG00000163513 | NM_001024847 | |

| | | | | | | | |
|---|---|---|---|---|---|---|---|
| TNFa | TNF | tumor necrosis factor | HGNC:11892 | P01375 | ENSG00000232810 | NM_000594 | [30, 33, 148, 225-227] |
| TRAF2 | TRAF2 | TNF receptor associated factor 2 | HGNC:12032 | Q12933 | ENSG00000127191 | NM_021138 | [18, 117, 228] |
| TSC | TSC1 | TSC complex subunit 1 | HGNC:12362 | Q92574 | ENSG00000165699 | NM_000368 | [63, 204, 229] |
| VEGF | VEGFA | vascular endothelial growth factor A | HGNC:12680 | P15692 | ENSG00000112715 | NM_001025366 | [31, 65, 197, 230, 231] |
| VIM | VIM | vimentin | HGNC:12692 | P08670 | ENSG00000026025 | NM_003380 | [62, 80, 112, 115, 116] |
| Wnt3 | WNT3 | Wnt family member 3 | HGNC:12782 | P56703 | ENSG00000108379 | NM_030753 | [19, 20, 24, 66, 69, 70, 73, 129, 232, 233] |
| ZEB1 | ZEB1 | zinc finger E-box binding homeobox 1 | HGNC:11642 | P37275 | ENSG00000148516 | NM_030751 | [40, 52, 56, 57, 114, 128, 145, 234-237] |
| ZEB2 | ZEB2 | zinc finger E-box binding homeobox 2 | HGNC:14881 | O60315 | ENSG00000169554 | NM_014795 | |
| ZO-1 | TJP1 | tight junction protein 1 | HGNC:11827 | Q07157 | ENSG00000104067 | NM_003257 | [5, 51, 55, 85, 109, 128] |
| ITGA5 | ITGA5 | integrin subunit alpha 5 | HGNC:6141 | P08648 | ENSG00000161638 | NM_002205 | [78, 124, 131, 178, 207, 238, 239] |
| ITGB1 | ITGB1 | integrin subunit beta 1 | HGNC:6153 | P05556 | ENSG00000150093 | NM_002211 | |
| ITGB3 | ITGB3 | integrin subunit beta 3 | HGNC:6156 | P05106 | ENSG00000259207 | NM_000212 | |
| c-FOS | FOS | Fos proto-oncogene, AP-1 transcription factor subunit | HGNC:3796 | P01100 | ENSG00000170345 | NM_005252 | [17, 106, 240, 241] |

| | | | | | | | |
|---|---|---|---|---|---|---|---|
| c-Jun | JUN | Jun proto-oncogene, AP-1 transcription factor subunit | HGNC:6204 | P05412 | ENSG00000177606 | NM_002228 | [14, 30, 49, 50, 225] |
| c-Myc | MYC | MYC proto-oncogene, bHLH transcription factor | HGNC:7553 | P01106 | ENSG00000136997 | NM_001354870 | [15, 47, 73, 79] |
| comp | COMP | cartilage oligomeric matrix protein | HGNC:2227 | P49747 | ENSG00000105664 | NM_000095 | [102, 242] |
| eiF4E | EIF4E | eukaryotic translation initiation factor 4E | HGNC:3287 | P06730 | ENSG00000151247 | NM_001968 | [54, 78] |
| p12 | CDK2AP1 | cyclin dependent kinase 2 associated protein 1 | HGNC:14002 | O14519 | ENSG00000111328 | NM_004642 | [243, 244] |
| p15 | CDKN2B | cyclin dependent kinase inhibitor 2B | HGNC:1788 | P42772 | ENSG00000147883 | NM_004936 | [45-47] |
| p16 | CDKN2A | cyclin dependent kinase inhibitor 2A | HGNC:1787 | P42771 | ENSG00000147889 | NM_000077 | [8, 15, 16, 245] |
| p21 | CDKN1A | cyclin dependent kinase inhibitor 1A | HGNC:1784 | P38936 | ENSG00000124762 | NM_078467 | [45, 49, 77, 245, 246] |
| p27 | CDKN1B | cyclin dependent kinase inhibitor 1B | HGNC:1785 | P46527 | ENSG00000111276 | NM_004064 | [77, 94, 247, 248] |
| p38 | MAPK14 | mitogen-activated protein kinase 14 | HGNC:6876 | Q16539 | ENSG00000112062 | NM_001315 | [4, 17, 29, 86, 153, 249] |
| p53 | TP53 | tumor protein p53 | HGNC:11998 | P04637 | ENSG00000141510 | NM_000546 | [53, 72, 245, 246, 250] |
| pRB | RB1 | RB transcriptional corepressor 1 | HGNC:9884 | P06400 | ENSG00000139687 | NM_000321 | [16, 45, 246] |
| ATB | LNCRNA-ATB | lncRNA activated by TGF-beta | HGNC:52657 | | | NR_160525 | [61, 71] |
| miR-124 | MIR124-1 | microRNA 124-1 | HGNC:31502 | MI0000443 | ENSG00000284321 | NR_029668 | [60, 120, 122, 251] |

| | | | | | | | |
|---|---|---|---|---|---|---|---|
| miR-1246 | MIR1246 | microRNA 1246 | HGNC:35312 | MI0006381 | ENSG00000283203 | NR_031648 | [70] |
| miR-1269 | MIR1269A | microRNA 1269a | HGNC:35337 | MI0006406 | ENSG00000221563 | NR_031673 | [252] |
| miR-192 | MIR192 | microRNA 192 | HGNC:31562 | MI0000234 | ENSG00000283926 | NR_029578 | [235] |
| miR-195 | MIR195 | microRNA 195 | HGNC:31566 | MI0000489 | ENSG00000284112 | NR_029712 | [253-256] |
| miR-200 | MIR200A | microRNA 200a | HGNC:31578 | MI0000737 | ENSG00000207607 | NR_029834 | [71, 112, 113, 145, 234, 257] |
| miR-24 | miR24-1 | microRNA 24-1 | HGNC:31607 | MI0000080 | ENSG00000284459 | NR_029496 | [258, 259] |
| miR-429 | MIR429 | microRNA 429 | HGNC:13784 | MI0001641 | ENSG00000198976 | ENSG00000198976 | [260-265] |
| miR-629 | MIR629 | microRNA 629 | HGNC:32885 | MI0003643 | ENSG00000207965 | NR_030714 | [266] |
| miR-942 | MIR942 | microRNA 942 | HGNC:33688 | MI0005767 | ENSG00000215930 | NR_030640 | [92, 267-269] |
| TPA | | Simple Molecule | | | | | [23, 270] |
| mTOR | mTOR1 | mTOR Complex 1 | HGNC:3942<br>HGNC: 30287<br>HGNC: 28611<br>HGNC: 24825 | P42345<br>Q8N122<br>Q6R327<br>Q9BVC4 | ENSG00000198793<br>ENSG00000141564<br>ENSG00000164327<br>ENSG00000167965 | NM_004958<br>NM_020761<br>NM_152756<br>NM_022372 | [18, 175, 271] |
| | mTOR2 | mTOR Complex 2 | | | | | |
